# An epigenetic clock for human skeletal muscle

**DOI:** 10.1101/821009

**Authors:** S Voisin, NR Harvey, LM Haupt, LR Griffiths, KJ Ashton, VG Coffey, TM Doering, JM Thompson, C Benedict, J Cedernaes, ME Lindholm, JM Craig, DS Rowlands, AP Sharples, S Horvath, N Eynon

**Affiliations:** Institute for Health and Sport (iHeS), Victoria University, Footscray, VIC 3011, Australia; Faculty of Health Sciences & Medicine, Bond University, Gold Coast, QLD 4226, Australia; Genomics Research Centre, Institute of Health and Biomedical Innovation, School of Biomedical Sciences, Queensland University of Technology, Brisbane, QLD 4000, Australia; Department of Neuroscience, Uppsala University, Sleep Research Laboratory, 751 23 Uppsala, Sweden; Department of Medical Sciences, Uppsala University, Uppsala SE-75124, Sweden; Department of Medicine, School of Medicine, Stanford University, Stanford, CA 94305, USA; Centre for Molecular and Medical Research, Deakin University, Geelong Waurn Ponds Campus, Locked Bag 20000, Geelong, VIC 3220, Australia; Epigenetics, Murdoch Children’s Research Institute, Royal Children’s Hospital, Parkville, VIC 3052, Australia; School of Sport, Exercise and Nutrition, Massey University, New Zealand; Department of Physical Performance, Norwegian School of Sport Sciences, Oslo, Norway; Stem Cells, Ageing & Molecular Physiology Unit, Exercise Metabolism and Adaptation Research Group, Research Institute for Sport and Exercise Sciences, Liverpool John Moores University, Liverpool, L3 3AF, UK; Department of Human Genetics and Biostatistics, David Geffen School of Medicine, University of California Los Angeles, Los Angeles, CA 90095, USA

**Keywords:** Skeletal muscle, epigenetic clock, aging, DNA methylation, epigenetic age, biological age

## Abstract

**Background:** Ageing is associated with DNA methylation changes in all human tissues, and epigenetic markers can estimate chronological age based on DNA methylation patterns across tissues. However, the construction of the original pan-tissue epigenetic clock did not include skeletal muscle samples, and hence exhibited a strong deviation between DNA methylation and chronological age in this tissue.

**Methods:** To address this, we developed a more accurate, muscle-specific epigenetic clock based on the genome-wide DNA methylation data of 682 skeletal muscle samples from 12 independent datasets (18-89 years old, 22% women, 99% Caucasian), all generated with Illumina HumanMethylation arrays (HM27, HM450 or HMEPIC). We also took advantage of the large number of samples to conduct an epigenome-wide association study (EWAS) of age-associated DNA methylation patterns in skeletal muscle.

**Results:** The newly developed clock uses 200 CpGs to estimate chronological age in skeletal muscle, 16 of which are in common with the 353 CpGs of the pan-tissue clock. The muscle clock outperformed the pan-tissue clock, with a median error of only 4.6 years across datasets (*vs* 13.1 years for the pan-tissue clock, p < 0.0001) and an average correlation of ρ = 0.62 between actual and predicted age across datasets (*vs* ρ = 0.51 for the pan-tissue clock). Lastly, we identified 180 differentially methylated regions (DMRs) with age in skeletal muscle at a False Discovery Rate < 0.005. However, Gene Set Enrichment Analysis did not reveal any enrichment for Gene Ontologies.

**Conclusions:** We have developed a muscle-specific epigenetic clock that predicts age with better accuracy than the pan-tissue clock. We implemented the muscle clock in an R package called *MEAT* available on Bioconductor to estimate epigenetic age in skeletal muscle samples. This clock may prove valuable in assessing the impact of environmental factors, such as exercise and diet, on muscle-specific biological ageing processes.

## Introduction

Ageing is the normal, progressive decline of function occurring at the cellular, tissue and organismal levels over the lifespan^1^. Ageing increases susceptibility to a wide range of diseases, including cardiovascular and neurodegenerative diseases, metabolic disorders and many cancers^1^. It is therefore important to identify early and potentially modifiable molecular mechanisms that occur with advancing age. Changes in epigenetic patterns constitute a primary hallmark of ageing in all tissues of the human body^2^. Epigenetic marks are cellular properties conferring the ability to remember a previous biological event^3^, and some of these marks are sensitive to environmental stimuli such as diet, sleep^4^ and exercise training^5,6^. Epigenetic changes with age are particularly well characterised at the DNA methylation level^7,8^, including skeletal muscle^9^.

The first DNA methylation-based estimator of chronological age (known as the pan-tissue *epigenetic clock*) was developed using a wide spectrum of tissues and nucleated cell types^7^. The resulting regression model could then be used to estimate the chronological age of tissue samples based on the DNA methylation levels of 353 CpGs (Cytosine-phosphate-Guanine dinucleotides). The difference between estimated DNA methylation age and chronological age reflects not only technical noise but also biologically meaningful variation seen in epidemiological studies linking epigenetic aging rates to mortality risk, Alzheimer’s disease, and many age-related conditions^10,11^. Age-related conditions are often associated with tissue-specific effects. For example, obesity is associated with strong epigenetic age acceleration in human liver samples but negligible effects in muscle tissue when assessed by the pan-tissue clock^12^.

Most tissues exhibit similar epigenetic ages according to the pan-tissue clock but there are a few exceptions. For example, the cerebellum has been found to age more slowly^13^. Conversely, female breast tissue exhibits an increased epigenetic aging rate, especially in younger women^7,14^. The construction of the original pan-tissue clock did not include any skeletal muscle samples; instead, the pan-tissue clock was tested on few skeletal muscle samples (n = 66) with a strong deviation observed between DNA methylation and chronological age in this tissue^7^. While the pan-tissue clock has many applications, tissue-specific clocks developed exclusively in a given tissue, provide higher accuracy and specific application to specific tissues. In particular, blood tissue provides the opportunity to develop accurate predictors of lifespan and healthspan^15,16^, which is particularly useful as blood samples are little invasive. Specific epigenetic clocks have been developed for fibroblasts, keratinocytes, and buccal swabs^17^, and for cord blood samples^18^. However, to the best of our knowledge, no study to date has tackled the challenge of developing an epigenetic clock that is specific to human skeletal muscle.

An epigenetic clock well calibrated in skeletal muscle could prove useful for studying the impact of environmental factors (e.g. exercise) on epigenetic ageing of this tissue, and the relationship with health and disease processes^19^. In general, skeletal muscle tissue is of great interest to aging researchers and clinicians because skeletal muscle mass is lost at a rate of 0.5–1% per year after age 50^20^. This muscle loss (sarcopenia) leads to a host of age related complications including frailty, as well as increased morbidity and mortality^21^. At the same time, skeletal muscle loses mitochondrial function and becomes increasingly resistant to insulin with age^22^. However, skeletal muscle is remarkably plastic, which makes it a highly responsive target tissue for lifestyle^22^. For example, changes in DNA methylation that occur with a healthy diet^23^, and exercise^5,6^ may be mechanistically involved in slowing down the ageing process^1^.

In the current study, we aimed to address the poor performance of the pan-tissue clock in muscle by developing a muscle-specific epigenetic clock. We hypothesise that by using a large number of human skeletal muscle DNA methylation profiles, we can develop a muscle-specific epigenetic clock that outperforms the pan-tissue clock and that can estimate chronological age with high accuracy. We utilised DNA methylation data to estimate epigenetic age in a total of 682 male and female skeletal muscle samples aged 18-89. We also conducted an epigenome-wide association study (EWAS) to discover genes whose methylation change with age in skeletal muscle. We have made the muscle clock freely available in an R package called *MEAT* (Muscle Epigenetic Age Test) on Bioconductor.

## Results

### Description of the 12 skeletal muscle DNA methylation datasets

We gathered skeletal muscle methylomes from 12 datasets generated with three different platforms: HM27, HM450 and the more recent HMEPIC, totalling n = 682 samples (Fig. 1, Additional file 1). Three datasets came from our own lab or collaborators, and the other nine were publicly available on the Gene Expression Omnibus (GEO) platform or the database of Genotypes and Phenotypes (dbGAP). Only three datasets included women and only two datasets included non-Caucasian individuals. Eight of the 12 datasets were paired designs (e.g. monozygotic twins discordant for disease or pre/post interventions, Additional file 1), meaning that some of the 682 muscle samples were taken from healthy individuals at baseline or after a control diet, while other samples were taken after an exercise intervention, a high-fat diet, sleep deprivation, insulin stimulation, or were from individuals with T2D. We chose to keep all samples in the development of the muscle clock, as none of these factors were associated with drastic changes in age acceleration. For details on each individual dataset such as sample collection and DNA methylation assay, see Additional file 2.

**Fig. 1.**
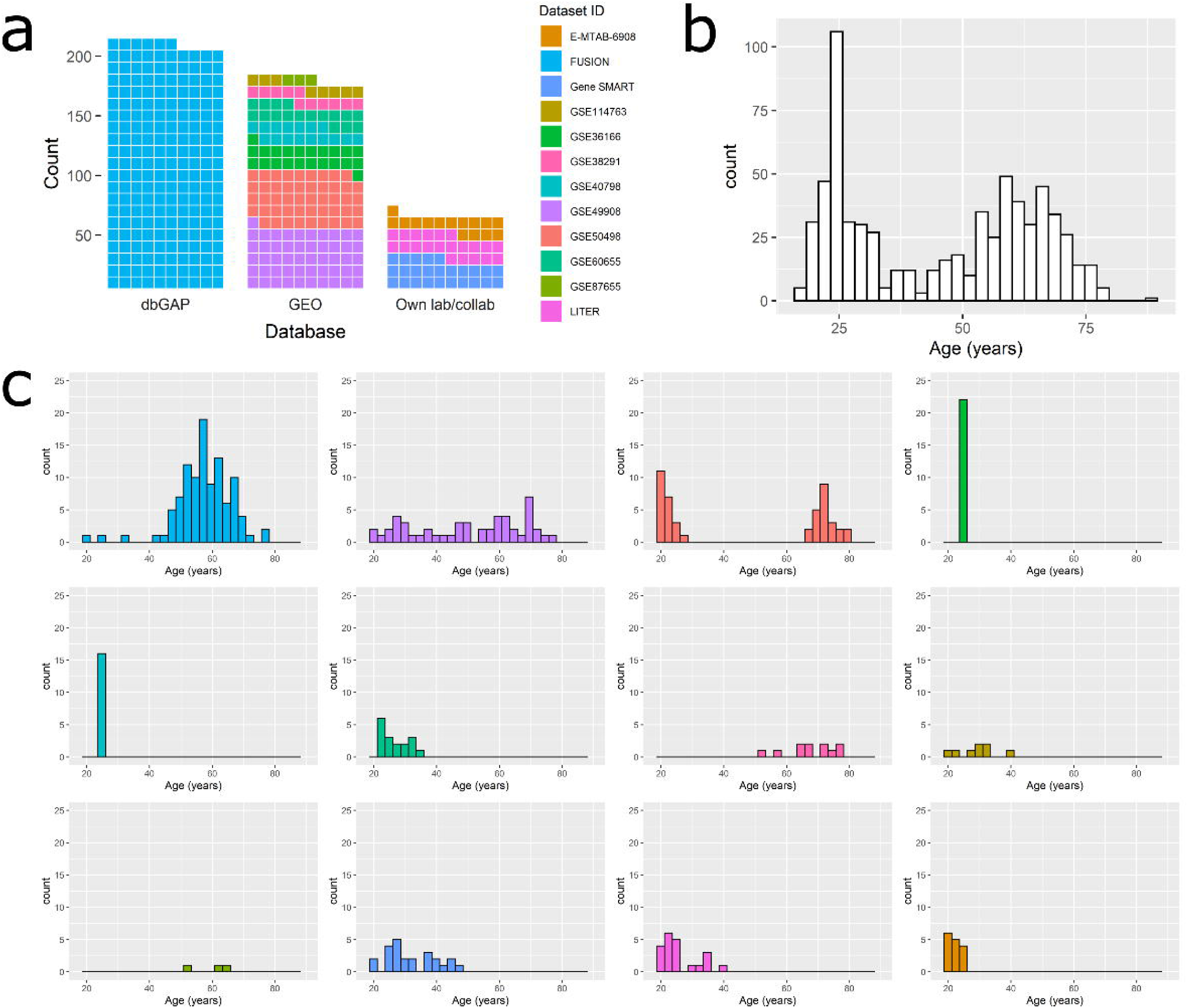
Age distribution in the 12 datasets used to develop the muscle clock. a) Waffle chart of the 12 datasets, split by database. Each cell represents 1 percentage point summing up to the total number of samples (n = 682); dbGAP = database of Genotypes and Phenotypes; GEO = Gene Expression Omnibus. b) Age distribution in all 12 datasets pooled together (n = 682). c) Age distribution in each individual dataset. Datasets were color-coded as in the waffle chart (a).

The 682 samples had a bimodal distribution of age, with an under-representation of 30-50 year olds (n = 242 aged 18-30, n = 105 aged 30-50, n = 275 aged 50-70, n = 60 aged 70-90, Fig. 1). More importantly, datasets greatly differed in their mean age and age range (Fig. 1, Additional file 1). For example, dataset GSE50498 contained younger (21.3 ± 2.4 years old), and older (73.2 ± 4.6 years old) but no middle-aged individuals; GSE36166 and GSE40798 had no variability in age, as all individuals were 24-25 years old.

### Development of a highly accurate skeletal muscle epigenetic clock

To develop the muscle clock, we adopted the same approach as Horvath^7^. Briefly, we restricted our analysis to the 19,401 CpGs that were present in all 12 datasets. Then, we used dataset GSE50498 that had a large sample size (n = 48), and the broadest age range (18-89 years old), as a gold standard to calibrate all other datasets. Although it does not entirely remove variability from different labs and platforms, this step allows for a harmonisation of DNA methylation profiles between datasets. Then, a transformed version of chronological age was regressed on the 19,401 CpGs using a penalized regression model (elastic net).

The elastic net model automatically selected 200 CpGs; with increasing age, 109 were hypomethylated and 91 hypermethylated (Additional file 3). Sixteen were in common with the 353 CpGs used in the pan-tissue clock (Fig. 2a). This is more than expected by chance, as none of the 1,000,000 randomly drawn samples of 200 CpGs from our dataset had more than 14 CpGs in common with the 353 CpGs of the pan-tissue clock. In addition, the effect of age on the methylation levels of 15/16 of these common CpGs was the same in both clocks. This shows that the muscle clock includes some CpGs whose methylation changes with age in all human tissues. We then tested for enrichment of the 200 muscle clock CpGs in CpG islands, and in skeletal muscle chromatin states. These chromatin states were determined by the Roadmap Epigenomics Project and provide a powerful, accurate mapping of gene and enhancer activity in human skeletal muscle at individual genomic positions. While we did not find any enrichment in CpG islands, shores or shelves, the muscle clock CpGs that were hypomethylated with age showed depletion in regions flanking active promoters (False Discovery Rate (FDR) = 0.00085, Fig. 2b).

**Fig. 2.**
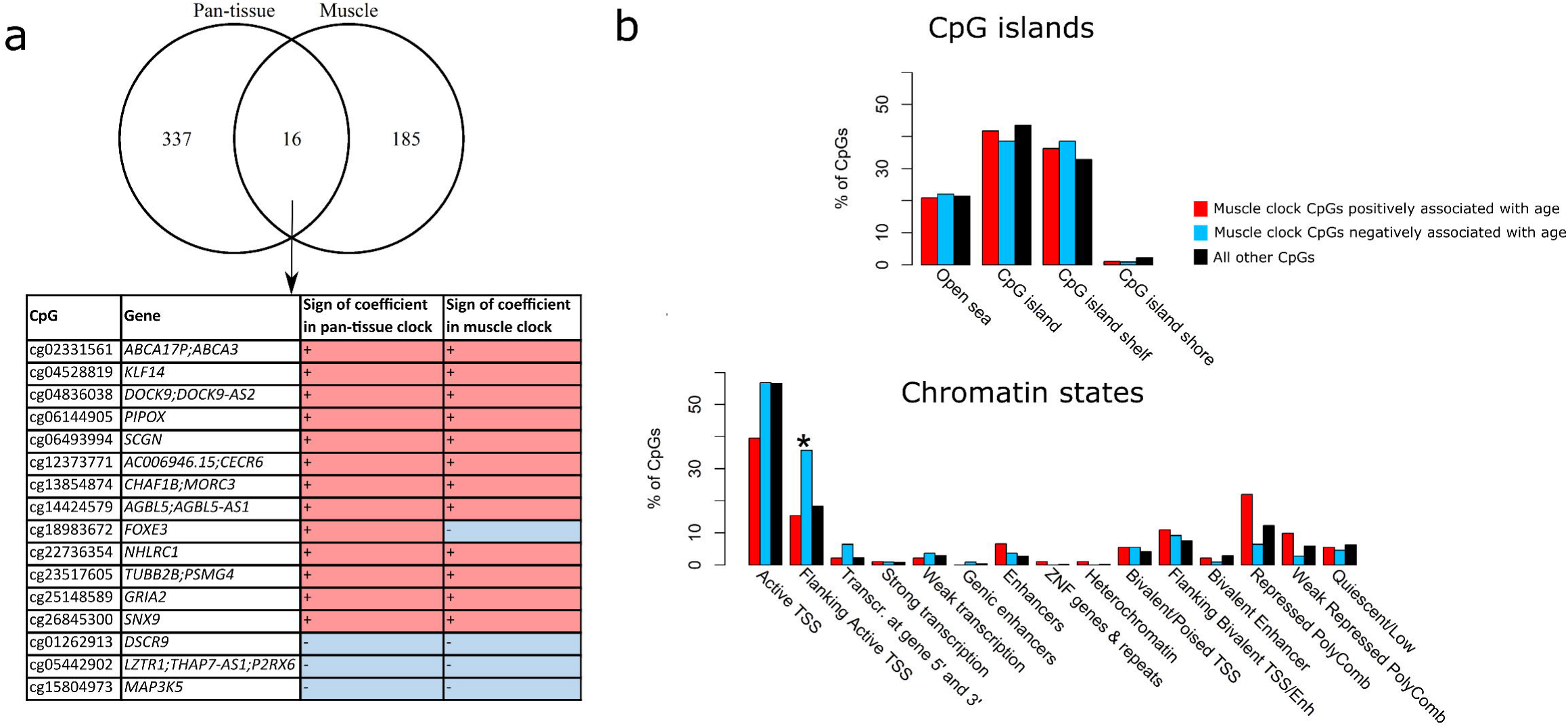
The 200 muscle clock CpGs. **a)** Overlap between the 353 CpGs of the pan-tissue clock and the 200 CpGs of the muscle clock. The 16 CpGs in common between the two clocks are displayed as table, with the annotated gene(s), and the direction of methylation with age in each of the two clocks. **b)** Enrichment of the 200 muscle clock CpGs in positions with respect to CpG islands (top), and in chromatin (bottom). Enrichment was tested with a Fisher’s exact test, adjusted for multiple testing. *FDR < 0.005.

### The muscle clock outperforms the pan-tissue clock

As the number of datasets and samples were rather limited (around six times fewer samples than those used to develop the pan-tissue clock), we adopted a leave-one dataset-out-cross-validation (LOOCV) procedure to obtain unbiased estimates of the muscle clock accuracy^7^. LOOCV is performed by removing one dataset and developing the clock on the 11 remaining datasets; the omitted dataset is then used as a test set (Fig. 3). Since we had 12 available datasets, we performed 12 LOOCVs (one for each dataset); this is better than performing a leave-one *sample*-out cross-validation procedure where the samples used to develop the clock contain samples from the same dataset as the omitted sample. This could lead to overly accurate age estimation, and would not apply well to new datasets. We then calculated three measures of accuracy: the correlation between predicted and actual age, the difference between predicted and actual age (AA_diff_), and the residual from a linear regression of predicted age against actual age (AA_resid_) (Fig. 3).

**Fig. 3.**
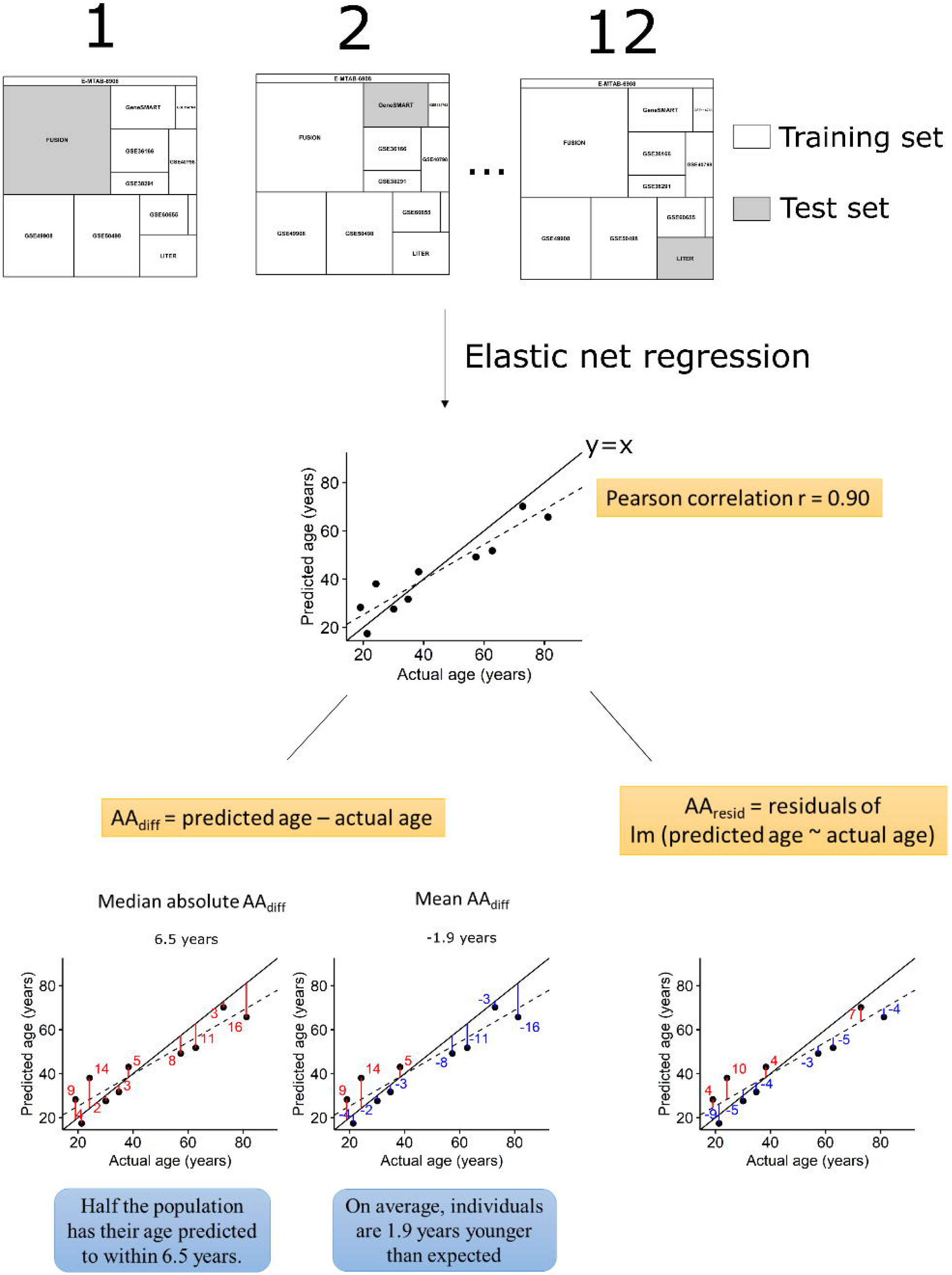
Methodology for leave-one dataset-out cross-validation (LOOCV) and measures of age prediction accuracy. In the LOOCV, one dataset is left out (test set) and all other datasets (training sets) are used to develop the age predictor. The DNA methylation profiles of the training sets are input into an elastic net regression model (*glmnet* package in R) and this model is then used to estimate age in the test set. Predicted and actual age were correlated using Pearson’s correlation coefficient (unless all individuals had the same age or the dataset was too small). We also calculated the AA_diff_ as the difference between predicted and actual age. We then calculated the median of the absolute values of AA_diff_ to estimate how well calibrated the clock was to this particular test set, and the mean of AA_diff_ to see whether the test set as a whole was younger (or older) than expected. Finally, we calculated the residuals from a linear regression of predicted age against actual age (AA_resid_) to obtain accuracy measures insensitive to the mean age of the dataset and to pre-processing techniques.

The skeletal muscle clock significantly outperformed the pan-tissue clock on the correlation between predicted and actual age (average ρ = 0.62 *vs* ρ = 0.51 across datasets, Fig. 4a, Additional file 4), and on the absolute AA_diff_ by 7.0 years (paired t-test p < 0.0001, Fig. 2b left panel, Additional file 4); however, the muscle clock was as accurate as the pan-tissue clock on the absolute AA_resid_ (paired t-test p = 0.16, Figure 2b right panel, Additional file 4). We also estimated the accuracy of the muscle clock by calculating the median absolute error and the average difference between predicted age and chronological age for each dataset ^7^. While the median absolute error is a robust measure of prediction error, the average difference indicates whether the predicted age of a given dataset is consistently higher (or lower) than expected^7^ (Fig. 3). Across the 12 datasets, the muscle clock performed very well, with a median absolute AA_diff_ of only 4.6 years on average (range 2.4-10.6 years) *vs* 12.0 years for the pan-tissue clock, and a median absolute AA_resid_ of 3.4 years on average *vs* 2.7 years for the pan-tissue clock (Additional file 4). Unsurprisingly considering the biased age distribution between and within datasets (Fig. 1, Additional file 1), both the muscle and pan-tissue clocks tended to predict younger ages for older individuals using AA_diff_ (Fig. 5). However, this bias was significantly reduced in the muscle clock, and was inexistent for AA_resid_ since by definition, AA_resid_ is unrelated to age.

**Fig. 4.**
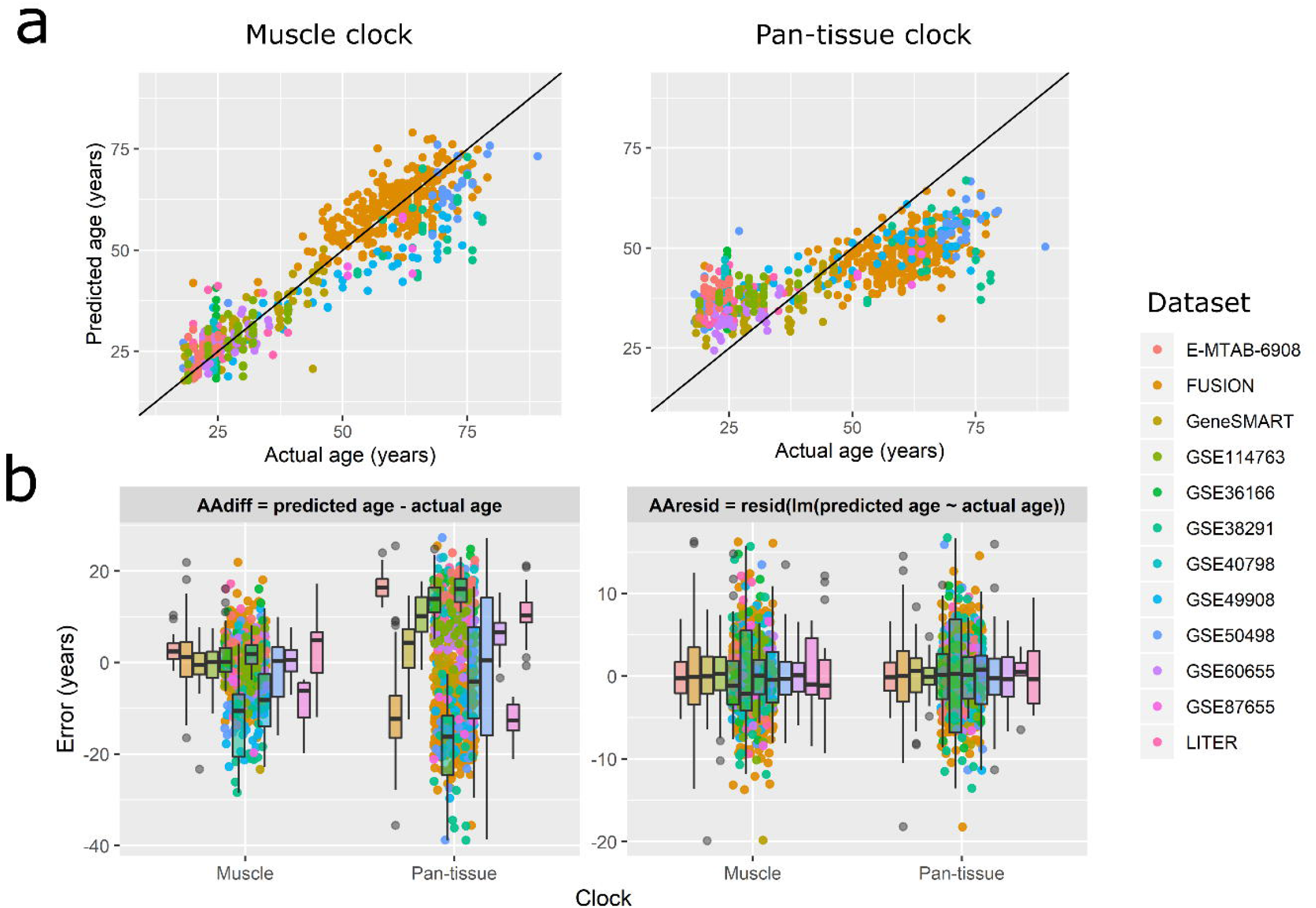
Predicted *vs* actual age and errors in age prediction in the LOOCV procedure. Each point corresponds to one of the 682 samples, colored by datasets to which they belong. a) Predicted *vs* actual age. Note that to obtain truly unbiased estimates of age prediction accuracy, the age predicted by the muscle clock is from the leave-one-out cross-validation procedure. b) Error in age prediction, either as the difference between predicted and actual age (left panel), or as the residuals from a linear model of predicted against actual age (right panel). Note that both panels are on different scales.

**Fig. 5.**
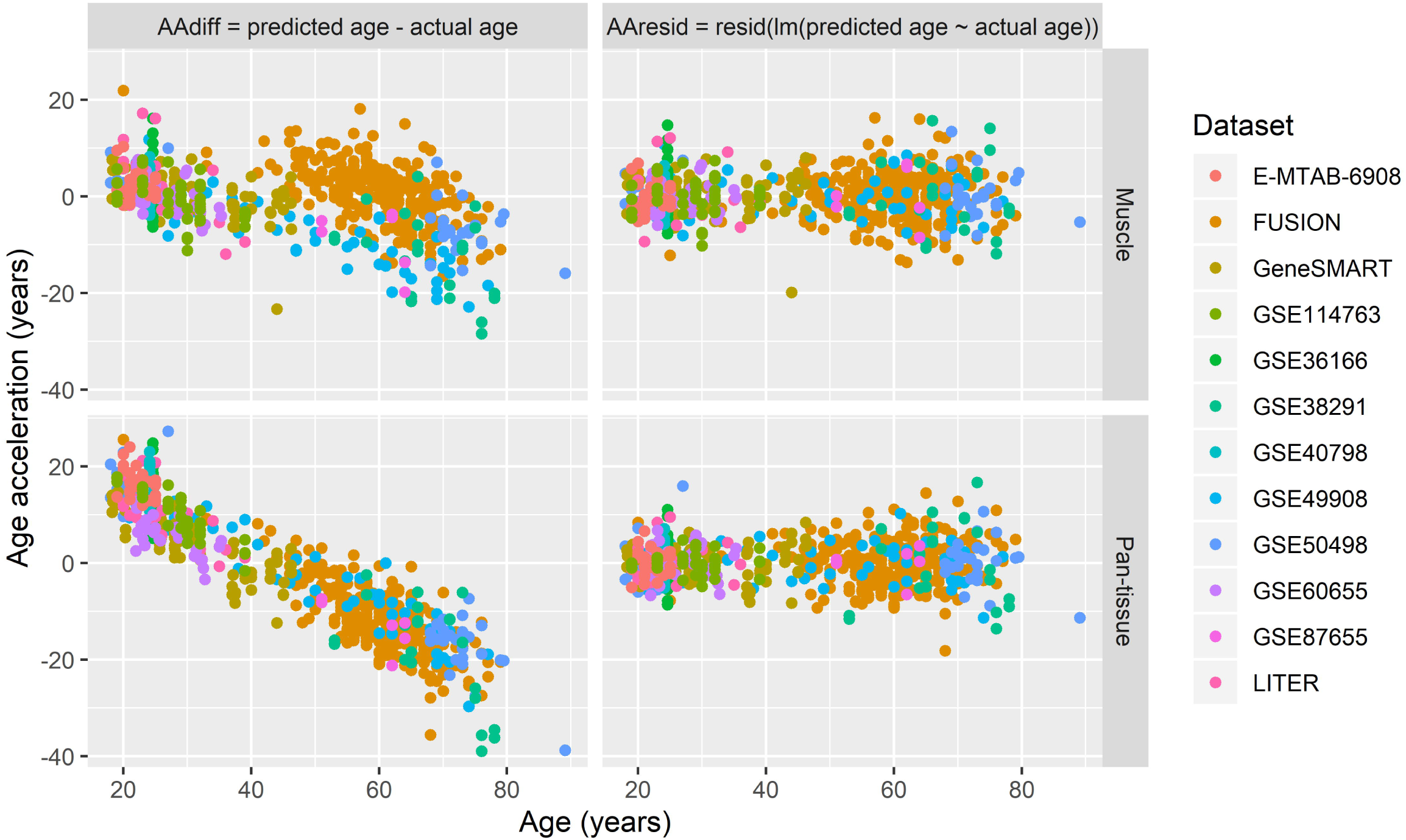
Age acceleration as a function of age in the muscle and the pan-tissue clocks. Here we show the bias in age prediction depending on the age of the individual. Using the difference between predicted and actual age (AA_diff_), younger individuals show systematically epigenetic ages than their real age, while older individuals show systematically older epigenetic ages than their real age. However, this bias is less pronounced in the muscle clock, and inexistent when using AA_resid_.

### EWAS of age

We took advantage of the large number of samples to explore DNA methylation patterns associated with age in skeletal muscle. We found 1,975 age-associated Differentially Methylated Positions (DMPs), corresponding to 180 Differentially Methylated Regions (DMRs) at FDR < 0.005 (Fig. 6a, Additional file 5). The direction of DNA methylation with age was balanced, with 51% of DMRs hypomethylated and 49% hypermethylated with advancing age (Additional file 5). 60% of the muscle clock CpGs were among the age-associated DMPs; one of these DMPs, located in Pipecolic Acid And Sarcosine Oxidase (*PIPOX*), was both in the muscle and pan-tissue clocks and showed one of the largest effect sizes (DNA methylation increased by 2.8% per decade of age, Fig 6b, Additional file 5). Both hypo- and hypermethylated DMPs were depleted in CpG islands and active TSS while simultaneously enriched in CpG island shelves, open sea, actively transcribed regions and enhancers (Fig. 6c). However, while hypomethylated DMPs were enriched in regions flanking active TSS and depleted in bivalent/poised TSS and in regions flanking bivalent TSS/enhancers, hypermethylated DMPs showed the opposite pattern (Fig. 6c). We then conducted a Gene Set Enrichment Analysis (GSEA) that takes into account the biased distribution of CpGs in genes, but found no enrichment of the DMPs for particular gene ontologies (GO) at FDR < 0.005.

**Fig. 6.**
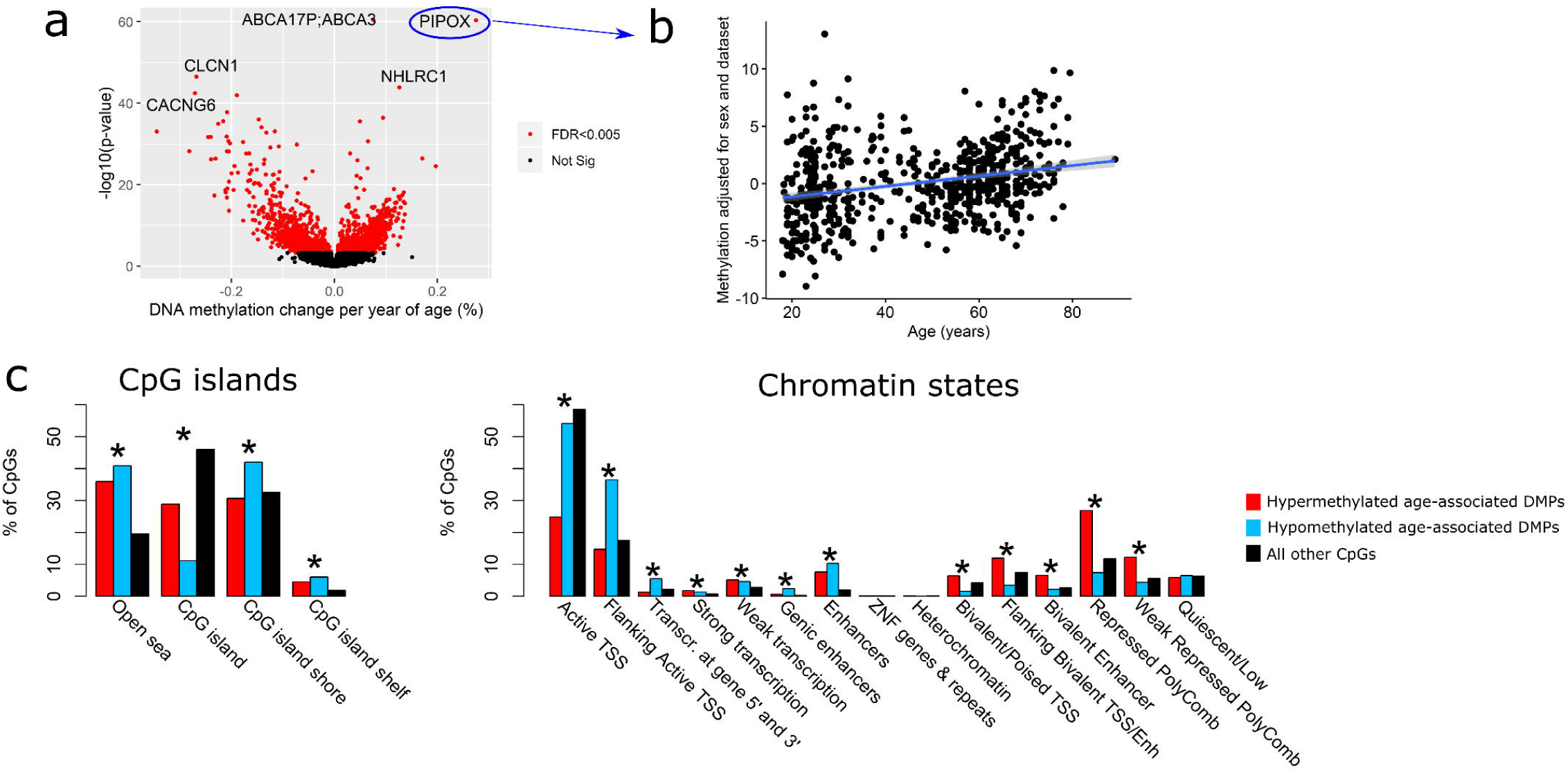
Differential DNA methylation with age in skeletal muscle. a) Volcano plot of DNA methylation changes with age. Each point represents a tested CpG (19,401 in total) and those highlighted in red were the differentially methylated positions (DMPs) significant at a false discovery rate (FDR) < 0.005. The x axis represents effect size, expressed as differential methylation per year of age. The y axis represents statistical significance, expressed as –log_10_(p-value), so CpGs that are higher on the graph are more significant. b) DNA methylation level as a function of age, for the CpG in *PIPOX* that was in both the muscle and pan-tissue clocks and that showed one of the largest effect size. c) Enrichment of DMPs in positions with respect to CpG islands (left), and in chromatin states (right). Enrichment was tested with a Fisher’s exact test, adjusted for multiple testing. *FDR < 0.005.

### *MEAT*: an R package to determine the epigenetic age of skeletal muscle

As part of the current investigation, we developed an open-access R package called *MEAT* (Muscle Epigenetic Age Test), available on Bioconductor. *MEAT* uses R code adapted from Horvath^7^ to calibrate skeletal muscle DNA methylation profiles to the GSE50498 gold standard. *MEAT* then calculates the epigenetic age of the calibrated samples using the muscle clock (elastic net model as implemented in *glmnet*). Users should provide a pre-processed β-value matrix generated with the Illumina HumanMethylation platform, as well as an optional phenotype table containing information such as age, sex, health/disease status, etc. If age is provided, the package will not only estimate epigenetic age, but also age acceleration (AA_diff_ and AA_resid_). Users can also ask *MEAT* to fit standard or robust linear models to test associations between phenotypes of interest (e.g. sex) and age acceleration in their datasets.

## Discussion

In the present study we developed an accurate epigenetic clock, specific to skeletal muscle, which outperformed the pan-tissue clock by an average of ∼7 years across 682 samples, in 12 independent datasets. This clock uses DNA methylation levels at 200 CpGs to predict chronological age, with a median absolute error of only 4.6 years, a significant improvement compared with the pan-tissue clock (12.0 years). We have made this clock available as an open-access R package called *MEAT* and available on Bioconductor. *MEAT* takes DNA methylation profiles assessed with the Illumina Infinium technology as input and outputs predicted age. This tool allows researchers to study the impact of environmental factors (e.g., exercise training, bed rest/immobilisation, diet, etc.) on the rate of aging in skeletal muscle samples. It could also be used to test whether diseased populations exhibit accelerated muscle-specific age acceleration compared with a matched healthy population, as was previously done using the pan-tissue clock^12,24–26^.

We highlighted some important limitations in age distribution both within and between datasets that could influence the accuracy of the muscle clock. Despite these limitations, the accuracy of the age predictor was excellent. The remarkable accuracy in prediction can be explained by multiple factors, most of which previously mentioned by Horvath^7^. First, the largest datasets (GSE50498, GSE49908, Gene SMART and FUSION) were also those with the broadest age range, which limits the confounding effect of age with dataset. Second, measurements from Illumina DNA methylation arrays are known to be less affected by normalization issues compared with those from gene expression (mRNA) arrays. Third, the elastic net model used to develop the epigenetic clock automatically selects CpGs that are less sensitive to differences in cohorts, labs and platforms since it is trained on datasets from various cohorts, labs and platforms. Fourth, the relatively large number of datasets helps average out spurious results and artefacts. Lastly, age affects DNA methylation levels of tens of thousands of CpGs^9^.

We found that there were more CpGs in common between the muscle- and pan-tissue clock^7^ than what would be expected by chance (as determined by our random sampling test). This suggests that the ageing process, despite being associated with many tissue-specific DNA methylation changes, is also associated with DNA methylation changes ubiquitous to all human cell types. The EWAS of age in skeletal muscle uncovered many genes whose methylation change with age. However, these genes were mostly distinct from the genes that are known to be differentially expressed in muscle with age^27^. Our relatively large sample size and wide age range allowed us to detect small effect sizes, and to uncover a large number of genes differentially methylated with aging in skeletal muscle. It is possible that age affects DNA methylation levels at these CpGs in all muscle cells. However, it is also possible that the DNA methylation differences between young and old individuals are due to differences in fibre type distribution, and perhaps also differences in satellite cell number and profiles. Slow- and fast-twitch fibres have distinct DNA methylation profiles^28^, and older muscle tends to have a greater proportion of slow-twitch fibres than young muscle^29^. In addition, satellite cells maintain their multipotent state via distinct DNA methylation profiles^30^ and both satellite cell numbers^31^ and DNA methylation profile^32^ change with age. The strength of this study lies in the utilisation of datasets that contained both young and older individuals from the general population; thus, it is likely that the muscle clock captured these DNA methylation changes due to fibre type changes with age. It was recently shown that controlling for heterogeneity in tissue/muscle fibre type reduces the number of physiological trait associations^33^, and it may also be the case that the epigenetic clock developed herein predicts different ages in different fibre types of a given individual. Uncovering which DNA methylation patterns change with age in fast-twitch fibres, slow-twitch fibres or in both fibres would be the next step to further enhance precision in the estimate of muscle age and in understanding how age affects muscle structure and function.

Skeletal muscle follows a circadian rhythm whose phase can be changed by environmental cues such as food, exercise and sleep^34^. Importantly, epigenetic mechanisms are involved in circadian rhythms and some DNA methylation oscillations were recently shown to happen at the same CpG sites that show age-related DNA methylation shifts in mice^35^. In the datasets we used to develop the muscle clock, most biopsies were taken in the morning in a fasted state, following a control diet for > 24h and exercise restriction for > 48h, which limits short-term environmental influences on DNA methylation levels. However, some datasets containing middle-aged and older individuals (GSE49908, GSE38291) did not have information on the conditions surrounding biopsy collection, so there is the possibility that some of these oscillations in DNA methylation are confounded with age in these datasets. We foresee that as more DNA methylation profiles in skeletal muscle are generated under controlled conditions and become publicly available, the muscle clock will be updated and gain in precision.

## Conclusions

In conclusion, we have developed an advanced muscle-specific epigenetic clock, using all known available datasets. This clock is freely available on Bioconductor as an R package (*MEAT*) for the scientific community to calculate epigenetic age. This new clock significantly outperforms the previous pan-tissue clock, and can calculate the epigenetic age in skeletal muscle with a mean accuracy of 4.9 ± 4.5 years across 682 samples. This muscle clock will be of interest and potential use to researchers, clinicians and forensic scientists working in the fields of skeletal muscle, chronic diseases, and ageing. In the future, we intend to evaluate how environmental factors, such as exercise and diet, could influence ageing via this newly developed clock.

## Methods

### Description of datasets used

We combined three datasets of DNA methylation in skeletal muscle (the Gene SMART (Skeletal Muscle Adaptive Response to Training)^36^, the E-MTAB-6908 study^4^, and the Bond University LITER study (unpublished)), with human skeletal muscle DNA methylation data from the open-access Gene Expression Omnibus (GEO) platform and the database of Genotypes and Phenotypes (dbGAP). We excluded datasets with < 3 samples, missing information on age (i.e. no age information on the GEO and corresponding author unresponsive), and datasets from primary cell culture experiments. Overall, we identified 8 datasets on the GEO and one dataset on dbGAP (Additional file 1), with sample sizes ranging from n = 3 to n = 282. Eight datasets were paired designs (e.g. monozygotic twins discordant for disease or pre/post interventions) and four cross-sectional. We described each dataset in details in the Supporting Information.

### Pre-processing

Whenever possible (i.e. when we had information on p-value detection for each probe, raw methylated and unmethylated signals or IDAT files, and batch/position information for each sample), we downloaded and pre-processed the raw data. If we did not have enough information on a given dataset (e.g. missing batch information), we utilised the processed data available on the GEO. In datasets that we did not pre-process, missing data was imputed using the champ.impute function of the *ChAMP* package^37^, with default parameters. As quality control, we ensured all datasets had a mean inter-correlation > 0.97 and a maximum beta-value > 0.99. For each individual dataset we pre-processed, we applied the following pre-processing steps using the R statistical software (www.r-project.org) together with the *ChAMP* analysis pipeline^37^ (for a full description of pre-processing steps on each dataset, see Additional file 1):

Any sample with > 10% of probes with detection p-value > 0.01 was removed (default parameter of the champ.load function). All probes with missing β-values, with a detection p-value > 0.01, probes with a bead count < 3 in at least 5% of samples, non-CG probes and probes aligning to multiple locations were removed, and for datasets containing males and females, probes located on the sex chromosomes were removed. SNP-related probes (“EUR” population probes in Zhou *et al*. ^38^) were also removed. β-values were obtained, and defined as

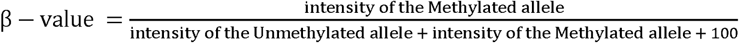

Then, β-mixture quantile normalization (BMIQ) method was applied to adjust for the Type I and Type II probe designs for methylation profiles generated from the HM450 and HMEPIC arrays. To identify technical and biological sources of variation in each individual dataset, Singular Value Decomposition (SVD) was performed. In all pre-processed datasets, both the plate and the position on the plate were identified as significant technical effects. Thus, all β-values were converted to M-values and the ComBat function from the *sva* package used to adjust directly for these technical artefacts.

Only 19,401 CpGs were identified to be in common between the 12 datasets after pre-processing, and all probes found on the HM27, HM450 and HMEPIC arrays (Additional file 6). To obtain DNA methylation profiles that were comparable between datasets, we adopted Horvath’s calibration method. We calibrated 11 or the 12 datasets to a gold standard, using the adapted version of the BMIQ algorithm^7^. We used GSE50498 as the gold standard since it was a large dataset (n = 48 samples) with the broadest age range (18-89 years old).

### Muscle clock development

We analysed 12 DNA methylation datasets from human skeletal muscle for which chronological age was available. We developed the muscle clock using an elastic net regression model identical to Horvath’s where a transformed version of chronological age was regressed on the 19,401 CpGs^7^. We first performed 10-fold cross-validation to select the optimal regularization parameter λ, using the elastic-net mixing parameter α = 0.5.

Given the limited number of datasets and the biased age distribution in each dataset, we adopted a leave-one-dataset-out cross-validation procedure to obtain an unbiased estimate of the muscle clock accuracy. We then calculated the prediction error as the age acceleration (AA), using two definitions that have been previously described^7,39^: the difference between predicted and actual age (AA_diff_), and the residual from a linear regression of predicted age against actual age (AA_resid_) (Fig. 3). While AA_diff_ is a straightforward way of calculating the error in age prediction, it is sensitive to the mean age of the dataset ^7^ and to the pre-processing of the DNA methylation dataset^39^; AA_diff_ can be biased upwards or downwards depending on how the dataset was normalized, and depending on the mean age and age variance of the dataset. In contrast, AA_resid_ is insensitive to the mean age of the dataset and is robust against different pre-processing methods^39^. Finally, we also calculated the Pearson correlation between predicted and actual age of the sample cohorts.

### Pan-tissue clock

We used the online epigenetic age calculator (https://dnamage.genetics.ucla.edu/home) selecting the option “Normalized Data” to implement the original pan-tissue clock^7^.

### Statistics

We used a paired t-test on the absolute AA (AA_diff_ or AA_resid_) to compare the accuracy of the muscle clock to that of the pan-tissue clock. As recently suggested to improve replicability in science^40^, a p-value < 0.005 was deemed significant.

To identify age-associated methylation positions (DMPs), we used linear models and moderated Bayesian statistics as implemented in the *limma* package^41^. The DNA methylation levels at 19,401 CpGs from the n = 682 muscle samples were regressed against age, sex, and dataset ID. We used the block design as implemented in lmFit to account for the paired designs of some datasets. DMPs associated with age at a false discovery rate (FDR) < 0.005 were deemed significant^40,42^. To identify differentially methylated regions (DMRs, i.e. clusters of DMPs with consistent DNA methylation change with age), we used the *dmrcate* package^43^.

To identify age-associated GO terms, we conducted a Gene Set Enrichment Analysis (GSEA) as implemented in the gometh function of the *missMethyl* package^44^, using our own improved annotation of the epigenome and largely based on Zhou *et al*.’s annotation^38^. This function accounts for the biased distribution of CpGs in genes. All GO terms pathways at FDR < 0.005 were deemed significant^40,42^.

To test whether the clock CpGs showed any enrichment inside or outside CpG islands, or enrichment for specific chromatin states, we compared the distribution of the clock CpGs with that of all other CpGs in different CpG island domains (open sea, CpG island, CpG island shore, CpG island shelf), or chromatin states in male skeletal muscle from the Roadmap Epigenomics Project with a Fisher’s exact test. As there are 4 different positions with respect to CpG islands and 15 different chromatin states, we only considered positions with respect to CpG islands and chromatin states significant if FDR < 0.005.

## Acknowledgements

This work was supported by Sarah Voisin’s National Health & Medical Research Council (NHMRC) Early Career Research Fellowship (APP11577321) and by Nir Eynon’s NHMRC Career Development Fellowship (APP1140644). The Gene SMART and LITER studies were both supported by the Collaborative Research Network for Advancing Exercise and Sports Science (201202) from the Department of Education and Training, Australia. Mr Nicholas Harvey was supported by a PhD stipend also provided by Bond University CRN-AESS.

This research was also supported by infrastructure purchased with Australian Government EIF Super Science Funds as part of the Therapeutic Innovation Australia – Queensland Node project (LRG). We also greatly acknowledge Erika Guzman at the ATGC/IHBI/QUT for performing the HMEPIC assays in the LITER study. We would also like to acknowledge Matthew McKenzie at Deakin University for coming up with the “*MEAT*” acronym. The authors adhere to the ethical guidelines for publishing in the journal of cachexia, sarcopenia and muscle^45^.

The authors declare that they have no conflict of interest.

## Additional files

**Additional file 1. Overview of the 12 datasets of DNA methylation in skeletal muscle**.

**Additional file 2. Detailed description of the 12 datasets of DNA methylation in skeletal muscle**.

**Additional file 3. Detailed information on the 200 CpGs automatically selected by the elastic net model.**

Coefficient = coefficient in the elastic net model. Each CpG was annotated to one or more genes using the annotation file from Zhou *et al*.^38^ to which we added annotation to long-range interaction promoters using chromatin states in male skeletal muscle from the Roadmap Epigenomics Project and GeneHancer information from the Genome Browser (hg38).

**Additional file 4. Leave-one dataset-out cross-validation (LOOCV) analysis of the muscle clock and comparison with the pan-tissue clock.**

Each row shows accuracy estimates for a given dataset. The three accuracy measures reported in this paper include the Pearson correlation coefficient between predicted and actual age, the difference between predicted age and actual age (AA_diff_), and the residuals from a linear model of predicted age against actual age (AA_resid_). Note, the shaded cells indicate we did not calculate the Pearson correlation coefficient for datasets GSE36166 and GSE40798 as they are invariant in age, nor for dataset GSE87655 as the sample size was too low (n = 3). Data are shown as mean ± SD.

**Additional file 5. Summary of differentially methylated positions and regions with age.**

Effect size = methylation change per year of age or between men and women. Each CpG was annotated to one or more genes using the annotation file from Zhou *et al*.^38^.

**Additional file 6. Genomic location and annotated genes for the 19**,**401 CpGs in common between all 12 datasets of DNA methylation in skeletal muscle.**

