## Additional file 2 for "An epigenetic clock for human skeletal muscle"

### Detailed information on the 12 DNA methylation datasets

##### **FUSION**

This dataset contained 282 *vastus lateralis* muscle samples from men and women aged 20-77 years old, collected in 3 different study sites in Finland^1^. ). DNA methylation profiles were generated with the Illumina HMEPIC array^2^. Participants had either normal glucose tolerance (NGT), impaired fasting glucose (IFG), impaired glucose tolerance (IGT) or type 2 diabetes (T2D).

##### **The Gene SMART (Skeletal Muscle Adaptive Response to Training) cohort**

This dataset contains 75 *vastus lateralis* muscle samples from healthy men who underwent an endurance training program^3^. From the 75 samples, 25 were taken at baseline, 25 after one bout of high-intensity interval exercise, and 25 after four weeks of exercise training (25 individuals at 3 time points). DNA methylation profiles were generated with the HMEPIC, yielding methylation levels at ~ 850,000 CpGs. The assay was performed in the Genomics Research Centre, Institute of Health and Biomedical Innovation, Queensland University of Technology, Brisbane, Australia. Total gDNA was extracted from 15 mg muscle tissue using the Allprep DNA/RNA/miRNA universal kit (Qiagen). The manufacturer’s protocol was modified to include the use of a Bullet Blender Gold Homogeniser (NextAdvance) in combination with dry ice for complete tissue digestion to ensure no enzymatic digestion of the DNA. Following isolation, DNA quantity and quality was measured using the dsDNA Broad Range Qubit kit (ThermoFisher scientific) on a Qubit Flourometer V2.0. For bisulphite conversion of gDNA for subsequent methylation arrays, 500 ng total gDNA was bisulfite converted using the EZ DNA methylation kit (ZYMO research) as per the manufacturer’s instructions. Bisulfite converted DNA (bDNA) was then used for the methylation array protocol. Briefly, bDNA was used for amplification of the regions of interest for binding to the Infinium BeadChip arrays. The amplification products were then fragmented, precipitated, resuspended, and then hybridised onto the BeadChips. The chips were washed and then stained prior to application on an HMEPIC array according to the manufacturer’s protocol. The HMEPIC arrays were then visualised and analysed using the Illumina HiScan instrument and GeneStudio software (Illumina).

##### **GSE49908**

This dataset contained 51 *vastus lateralis* muscle samples from healthy men aged 21-77 years old, collected at the Department of Physiology and Biophysics and the Centre for Aging at the University of Alabama at Birmingham^4^. DNA methylation profiles were generated with the Illumina HM27 array.

##### **GSE50498**

This dataset contained 48 *vastus lateralis* muscle samples from healthy younger and older men^5^. DNA methylation profiles were generated with the Illumina HM450 array.

##### **GSE36166**

This dataset contained 44 *vastus lateralis* muscle samples from healthy men of the same age (24.6 ± 1.1 years old)^6^. From the 44 samples, 22 were taken after a control diet and the other 22 following a high-fat diet (21 paired samples). DNA methylation profiles were generated with the Illumina HM27 array.

##### **LITER (Limb Immobilisation and Transcriptional/Epigenetic Responses) cohort**

This dataset contained 21 *vastus lateralis* muscle samples from healthy men aged 20-39 prior to a 14-day limb immobilisation program (Australia New Zealand Clinical Trials Registry: ACTRN12616001399482). DNA methylation profiles were generated with the HMEPIC array as described for The Gene SMART cohort, except as follows: total gDNA was extracted from 35 mg muscle tissue using the Allprep DNA/RNA/miRNA universal kit (Qiagen). The HMEPIC array assays were performed at the Australian Translational Genomics Centre, Institute of Health and Biomedical Innovation, Queensland University of Technology, Brisbane, Australia.

##### **GSE40798**

This dataset contained 31 *vastus lateralis* muscle samples from healthy men with similar age (24.1 ± 0.5 years old), with a low birth weight^7^. From the 31 samples, 16 were taken after a control diet and the other 15 after a high-fat diet (15 paired samples). DNA methylation profiles were generated with the Illumina HM27 array.

##### **GSE60655**

This dataset contained 34 *vastus lateralis* muscle samples from healthy men and women aged 21-35 years old who underwent a 12-week long endurance training program^8^. From the 34 samples, 17 were taken at baseline and 17 after the intervention (17 paired samples). DNA methylation profiles were generated with the Illumina HM450 array.

##### **E-MTAB-6908**

This dataset contained 30 *vastus lateralis* muscle samples from 15 healthy men aged 19-25 who took part in two conditions (overnight wakefulness vs sleep)^9^. DNA methylation profiles were generated with the Illumina HM450 array.

##### **GSE38291**

This dataset contained 22 *vastus lateralis* muscle samples from 11 pairs of monozygotic twins aged 53-80 years old and discordant for Type 2 Diabetes^10^. DNA methylation profiles were generated with the Illumina HM27 array.

##### **GSE114763**

This dataset contained 40 *vastus lateralis* muscle samples from healthy men aged 19-39 years old, who underwent a resistance training program twice, with a washout period in between^11^. From the 38 samples, 8 were taken at baseline, 8 after an acute (one session) bout of resistance exercise, 8 after the first intervention, 8 after 7 weeks of washout, and 8 after the second intervention (8 individuals at 5 time points). DNA methylation profiles were generated with the Illumina HMEPIC array.

##### **GSE87655**

This dataset contained 6 *vastus lateralis* muscle samples from healthy men aged 51-62, incubated in either a control buffer or insulin^12^. Out of the 6 samples, 3 were incubated with control buffer and the other 3 incubated with insulin (3 paired samples). DNA methylation profiles were generated with the Illumina HM450 array.

##### **GSE58280**

This dataset contained 34 *vastus lateralis* muscle samples from middle-aged Pacific Islanders who underwent a 16-week long endurance or resistance training program^13^. Out of the 34 samples, 17 were taken before and the other 17 after the intervention (17 paired samples). DNA methylation profiles were generated with the Illumina HM450 array. We initially intended to use this dataset as a test set, but in our initial screening, predicted sex based on DNA methylation levels did not match the assigned sex of the samples. Therefore, this dataset was excluded from further analysis.
